# *De novo* evolution of transmissible tumors in Hydra

**DOI:** 10.1101/2024.03.13.584762

**Authors:** Sophie Tissot, Jordan Meliani, Justine Boutry, Lionel Brazier, Jácint Tökölyi, Benjamin Roche, Beata Ujvari, Aurora M. Nedelcu, Frédéric Thomas, Antoine M. Dujon

## Abstract

While most cancers are not transmissible, there are rare cases where cancer cells have acquired the ability to spread vertically or horizontally to other individuals, and sometimes species, causing epidemics in their hosts. However, as these transmissible cancers are usually detected once they are relatively well disseminated in host populations, the conditions associated with their origin remain poorly understood. Using the freshwater cnidarian *Hydra oligactis*, which exhibits spontaneous tumor development that in some strains became vertically transmitted, this study presents the first experimental observation of the evolution of a transmissible tumor. Specifically, we assessed the initial vertical transmission rate of spontaneous tumors and explored the potential for optimizing this rate through artificial selection. One of the hydra strains, which evolved transmissible tumors over five generations, was characterized by analysis of cell type and microbiome, as well as assessment of life-history traits. Our findings indicate that tumor transmission can be immediate for some strains and can be enhanced by selection. The resulting tumors are characterized by overproliferation of large interstitial stem cells and, in contrast with other transmissible tumors on Hydra, are not associated with a specific microbiome. Furthermore, although tumor transmission has only been established over 5 generations, it was sufficient to alter life-history traits in the host, suggesting a compensatory response. This work, therefore, makes the first contribution to understanding the conditions of transmissible cancer emergence and their short-term consequences for the host.

## Introduction

The evolution of animal multicellularity at the end of the Precambrian was accompanied by the ability to regulate cell proliferation in a spatio-temporal context. However, mutations in these regulatory systems can occur, leading to uncontrolled proliferation that can result in neoplasms (tumors) that, in some cases, transform into invasive and deadly cancers [1][2]. Even though malignant progressions can and do occur in the majority of multicellular organisms, only a few cases of transmissible cancers have been reported (although their number could be underestimated, see [3])[4]. Currently, fourteen transmissible cancers are known in the wild: two affecting the Tasmanian devil [5], one observed in the Canidae [6], and eleven identified in bivalves (in some of the latter cases, they also exhibit the capacity to cross the species barrier) [7], [8], [9], [10], [11]. The transmissibility of these cancer cells aligns their evolutionary dynamics more closely with those of emerging pathogens, fostering long-term co-evolution with their host. For example, about forty years after its manifestation [12], the transmissible cancer line associated with the Devil Facial Tumor Disease (DFTD) in Tasmanian devils seems to evolve into an obligatory parasite that genuinely co-evolves with its host [13].

Transmissible tumor cells, in addition to expressing the hallmarks of cancer cells [14], have to match a series of exact conditions (i.e., a “perfect storm”) to emerge and spread as an epidemic. According to Ujvari et al. [15], the perfect storm theory considers at least four key factors, namely (i) the release of tumor cells from the impacted host, (ii) the persistence of tumor cells during transit between hosts, (iii) a conducive environment supporting invasion, and (iv) the ability to adapt to new environments while avoiding immune responses in the foreign host. In 2022, Tissot et al. [16] uncovered an additional hurdle to the emergence of transmissible cancer, beyond acquiring transmissibility itself: the ability for dissemination within host populations, at least until a prevalence threshold is reached, triggering the epidemic. The key parameters influencing the crossing of this second barrier pertain to abiotic or biotic variables that act directly on the cancer cells (e.g. water current in bivalves [17]) and/or on the infected hosts (e.g., predation, parasitism, [15][17][18][19][20]), potentially halting the outbreak of an epidemic even if the first transmissibility barrier has been breached. The many conditions required to create a perfect storm could explain the rarity of transmissible cancers [16]. Furthermore, a significant limitation to our understanding of the conditions leading to the emergence of transmissible cancers is that their presence is typically observed only after they have spread widely within host populations, i.e., relatively long after their initial appearance (e.g. [12]).

The freshwater cnidarian *Hydra oligactis* has been observed to spontaneously develop tumors under laboratory conditions, particularly in response to extensive feeding [21][22]. In a notable case that occurred 15 years ago in Thomas Bosch’s laboratory in Germany, a hydra developed a tumor, consisting of overproliferation of large interstitial stem cells, capable of being transmitted vertically through the asexual reproduction of its host, by budding. The isolation and culture of this polyp enabled the establishment of a tumoral hydra line, referred to as the St. Petersburg strain [24]. As the bud detaches from the tumoral hydra, it undergoes an initial prepathological phase, during which the tumors have not yet developed to the point of being detectable externally. However, after approximately four or five weeks, tumors become visible, marking the transition into the pathological phase [24]. Subsequent research revealed that some of these transmissible tumors were linked to a specific microbiome, notably involving the co-occurrence of spirochetes and *Pseudomonas* [25]. Additionally, Boutry et al. [23] recently demonstrated that some wild *H. oligactis*, when brought back to the laboratory, develop spontaneous tumors at relatively high frequencies (i.e. up to 30%, see [22]), depending on the population of origin. However, it remains unclear to what extent these spontaneous tumors are already transmissible, and/or if this trait can be acquired over time. In this context, the hydra-tumor model appears to be an excellent model for attempting to explore, for the first time, the evolution of transmissible tumors.

In this study, employing *H. oligactis*, we observed for the first time the experimental evolution of a transmissible tumor. Our aim was to assess the initial transmission rate of these tumors and explore the potential for optimizing this rate through selection. One of the selected strains was then examined through cell type analysis, and its microbiome as well as life history traits were explored.

## Results

### Selection for tumor transmissibility

To assess the conditions associated with the emergence and evolution of transmissible tumors in *H. oligactis*, we investigated the presence of tumors in the offspring (F1) developed from buds of wild individuals that had developed spontaneous tumors (F0). For those that had developed tumors, their offspring were also investigated over three additional generations. The results of this experimental selection are summarized in Figure 1. To control for the rate of spontaneous tumor appearance, buds from tumor-free hydra from the same wild population were surveyed in the same way.

**Figure 1.**
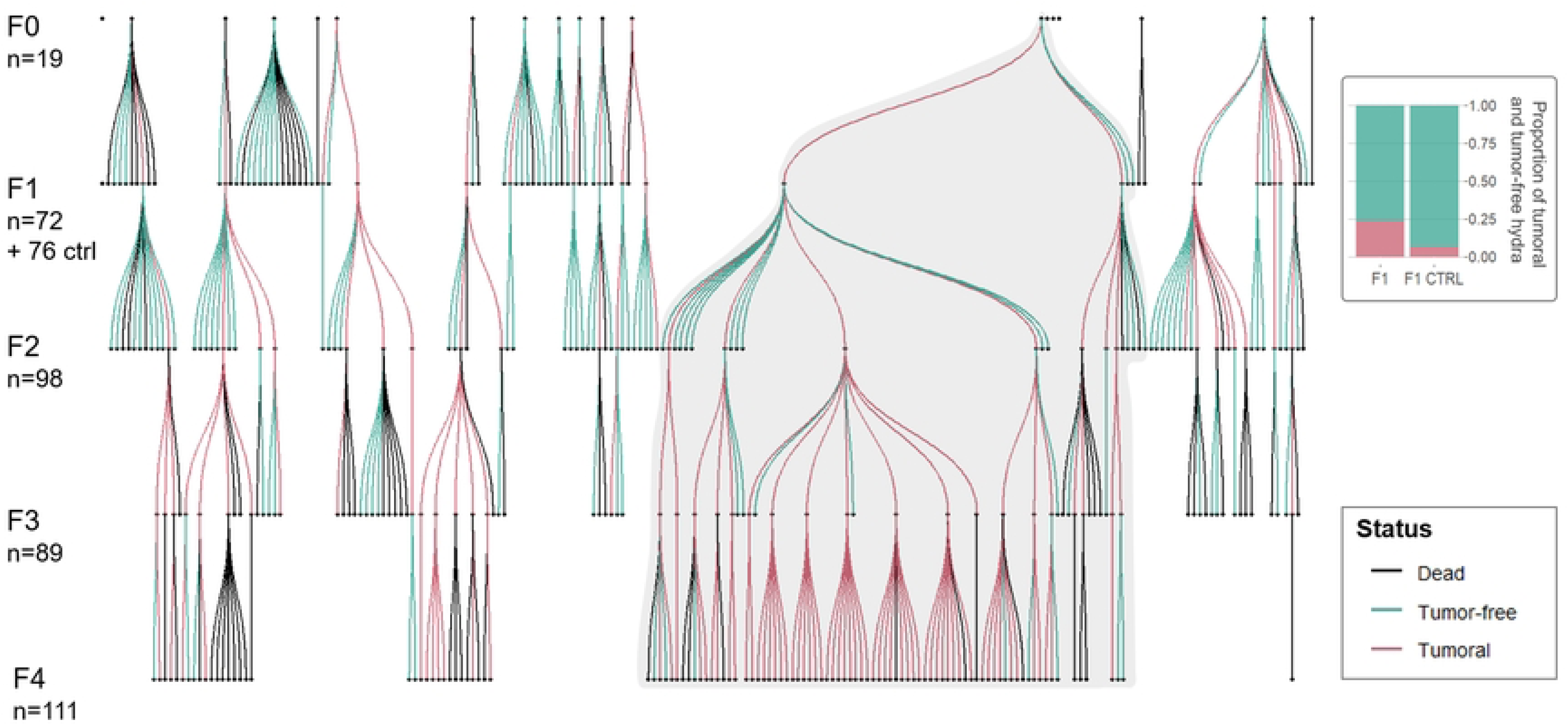
Summary of tumor transmission selection results and the proportion of hydras developing tumors in F1 and F1 CTRL. The MT40 strain, used for life-history trait analysis in the second part, is shaded.

#### Prevalence of tumors in the first generation

In the first generation, F1 hydras were about four times more likely to develop tumors when they came from tumoral F0 parents than from tumor-free parents (Figure 1; GLMM, OR = 4.27, SE = 2.30, p-value = 0.007). In the tumor-free population, about 7% (i.e. 5 out of 74) of individuals developed tumors, while in the tumoral strain, about 24% (i.e. 17 out of 72) developed tumors.

#### Estimation of transmission rate over generations

Concerning the tumor transmission rate, the model selected included the additive effect between generations and bud order, with a random effect of the founder individual. The tumor transmission rate more than doubled over the course of the experiment, from 35% in the first generation to 84% in the fourth (Figure 2; GLMM, *generation effect*, OR = 2.13, SE = 0.37, p-value < 0.001). The order of the bud formation has no significant effect on the transmission rate. Thus, the tumor transmission rate responded positively to artificial selection, but its occurrence among buds seemed to be random within an individual line.

**Figure 2:**
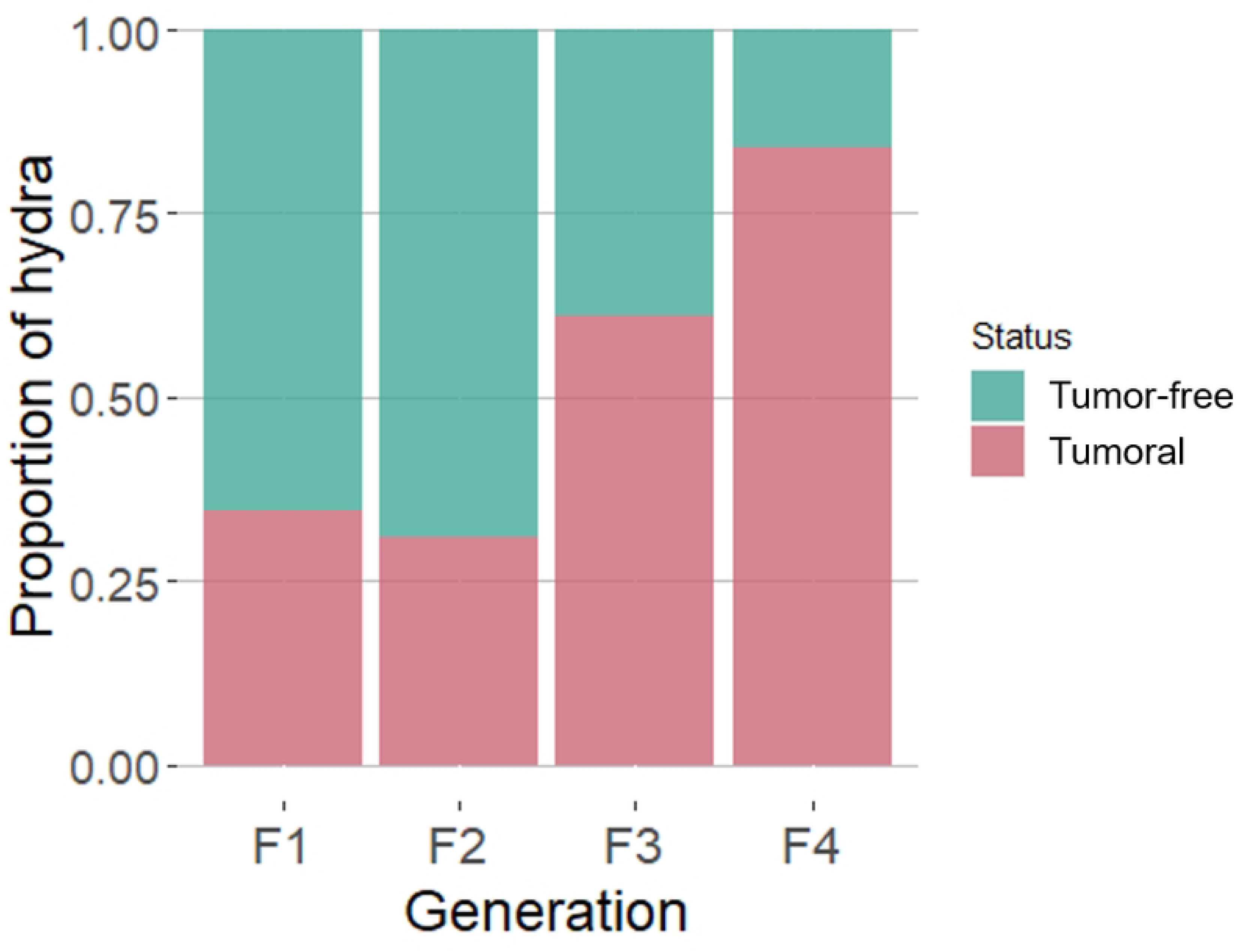
Proportion of hydras developing tumors according to generation.

### Life-history traits

Using hydras from the tumoral strain MT40 (established during the previous tumor transmission selection), and tumor-free wild hydras as control, we analyzed (i) age at first budding, (ii) asexual reproduction effort, (iii) bud survival, (iv) transmission rate and (v) survival time, according to the status (i.e. tumor-free, tumoral, and TFTP) and the phase (i.e. weeks 0 to 4, and weeks 5 to 9 matching for tumoral hydra to the prepathological and pathological phases) when relevant.

The status of the hydra had no effect on the age at first budding, as indicated by the model selection. On average, the first budding occurred around 17 days regardless of whether the hydra was tumoral or not.

The budding rate was explained by the interaction between the status of hydras and the phase with a random effect of the individual. Precisely, before 5 weeks, tumor-free hydras produced around 30% less buds than tumoral hydras and TFTP (Figure 3; GLMM, *status effect*, IRR = 0.68, SE = 0.09, p-value = 0.002). However, tumoral hydras were the only ones for which a decrease in the budding rate was observed after the fifth week, which corresponds to the start of the pathological phase for these hydras (Figure 3; GLMM, *interaction effect of status and phase*, IRR = 0.61, SE = 0.13, p-value = 0.024). The budding rate of tumoral hydra in pathological phase was reduced by about 40% compared to the prepathological phase.

**Figure 3.**
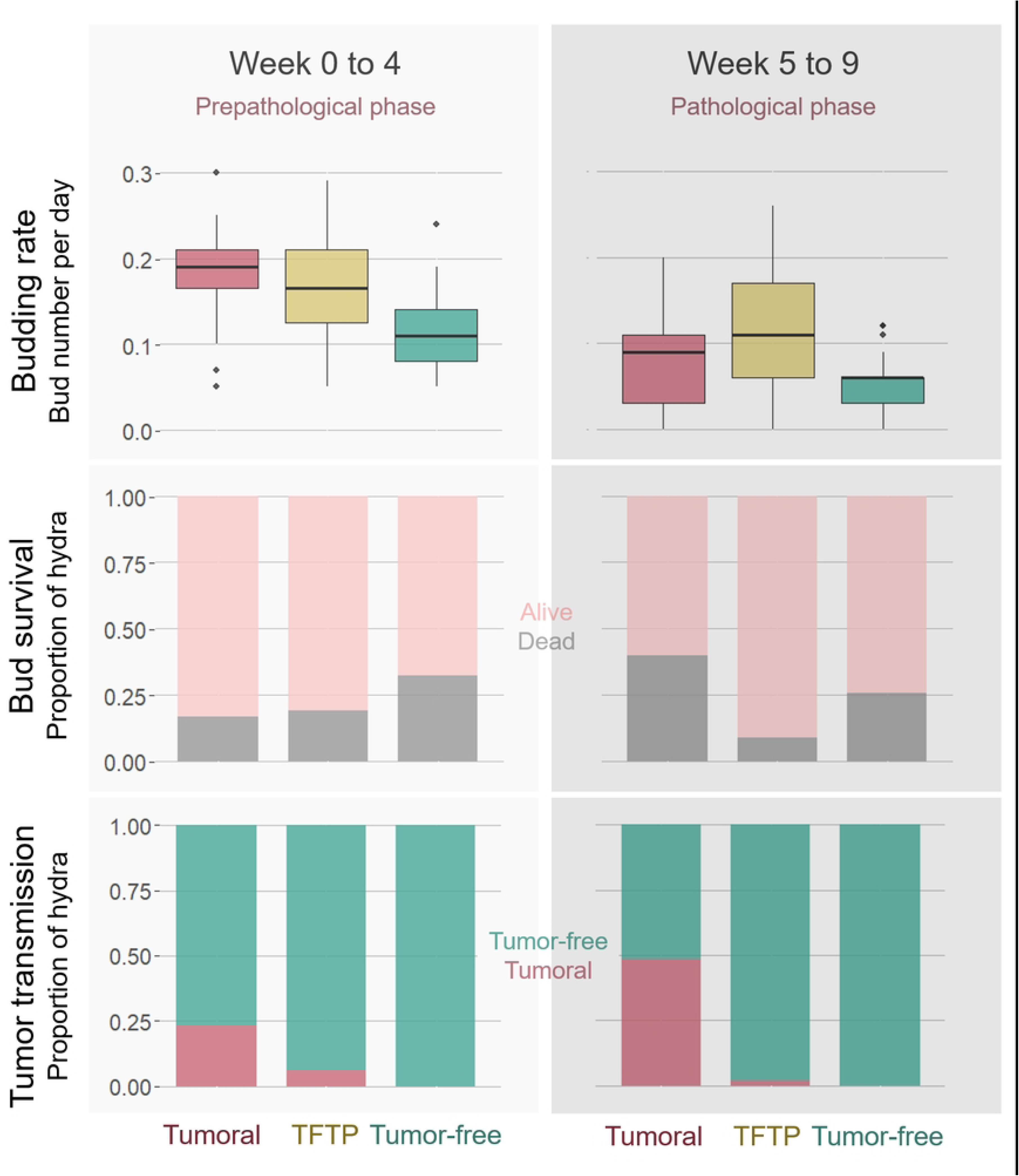
Budding rate, bud survival within two months, and tumor transmission rate according to hydra status for each phase.

For bud survival, the interaction effect between the phase in which the parents were at the time of bud production and the status were retained, as well as the random effect of parent identity. Buds of the three statuses showed a similar mortality risk, around 25%, when they are produced during the weeks 0 to 4. Only buds from tumoral hydras showed an increased mortality risk about 25% (from 15% to 40%) when they were produced during the weeks 5 to 9 corresponding for parents to the pathological phase, compared to weeks 0 to 4 where tumors are still unnoticeable (Figure 3; GLMM, OR = 8.10, SE = 5.88, p-value = 0.004). For buds issued from tumor-free hydras and TFTP, the mortality risk remained the same between the two phases.

The analysis of tumor transmission rate was conducted only on buds from tumoral hydras as no tumors were observed on buds from tumor-free control hydras. Thus, the model predicting the transmission rate in the MT40 tumoral strain included the parental status and the phase as fixed effects and controlled for the parental identity as a random effect. The chances of developing tumors are greatly reduced when the bud is derived from TFTP (Figure 3; GLMM, *effect of the parental status*, reference group: tumoral hydras, OR = 0.02, SE = 0.02, p-value = 0.001). On the other hand, when the bud was produced by a parent which subsequently developed a tumor, its probability of becoming tumoral was about four times higher. This is also the case if it is produced during the pathological phase of the parent, i.e. when the parent has already developed a tumor (Figure 3; GLMM, *effect of the phase of bud production*, reference group: week 5 to 9, OR = 0.36, SE = 0.19, p-value = 0.049).

Regarding survival time, tumoral hydras, tumor-free hydras and TFTP all have a half-life of around 100 days, as no effect was selected during model selection.

### Cell type and microbiome

Concerning to the cell type composition presented in Table 1, tumoral hydras showed a tenfold higher ratio of large interstitial stem cells to epithelial cells compared to TFTP and tumor-free hydras. In addition, their large to small interstitial stem cells ratio was 20 times higher than that observed in the other two statuses. Thus, as the ratio of small interstitial stem cells to epithelial cells was similar for all three statuses, the tumors seemed to result from overproliferation of large interstitial stem cells.

**Table 1.** Cell type composition for each status.

| Status | Ratios |  |  |
| --- | --- | --- | --- |
|  | LISC/EC | SISC/EC | LISC/SISC |
| Tumoral | 2.25 | 0.63 | 3.56 |
| TFTP | 0.15 | 0.87 | 0.18 |
| Tumor-free | 0.25 | 1.39 | 0.18 |

Ratios of large interstitial stem cells to epithelial cells (LISC/EC), small interstitial stem cells to epithelial cells (SISC/EC), and large interstitial stem cells to small interstitial stem cells (LISC/SISC) according to the status.

Regarding the abundance of the ten most represented order of bacteria in tumoral hydras, tumor-free and TFTP, they all appear to be mainly colonized by Chlamidiales (see Figure 4). Alpha diversity did not differ significantly between the three groups (Kruskal-Wallis χ^2^ = 1.82, p = 0.403). However, the Adonis analyzes showed that the microbiome composition of the three groups differed from each other significantly or marginally significantly (Bray-Curtis index: F = 1.97, p = 0.054; Jaccard presence-absence index: F = 1.536, p = 0.002). Nevertheless, in both cases it was the tumor-free group that differed from the other two (tumor-free vs. TFTP, Bray-Curtis index, adjusted p = 0.032; Jaccard presence-absence index, adjusted p = 0.024; tumor-free vs tumoral, Bray-Curtis index, adjusted p = 0.032; Jaccard presence-absence index, adjusted p = 0.095), while tumoral and TFTP showed no significant differences (tumoral vs. TFTP, Bray-Curtis index, adjusted p = 0.286; Jaccard presence-absence index, adjusted p = 0.114). The principal coordinate analyses of the Bray-Curtis index and Jaccard index according to the status are available in the supporting information (S1 Figure), along with a graphical representation of alpha diversity for each status.

**Figure 4:**
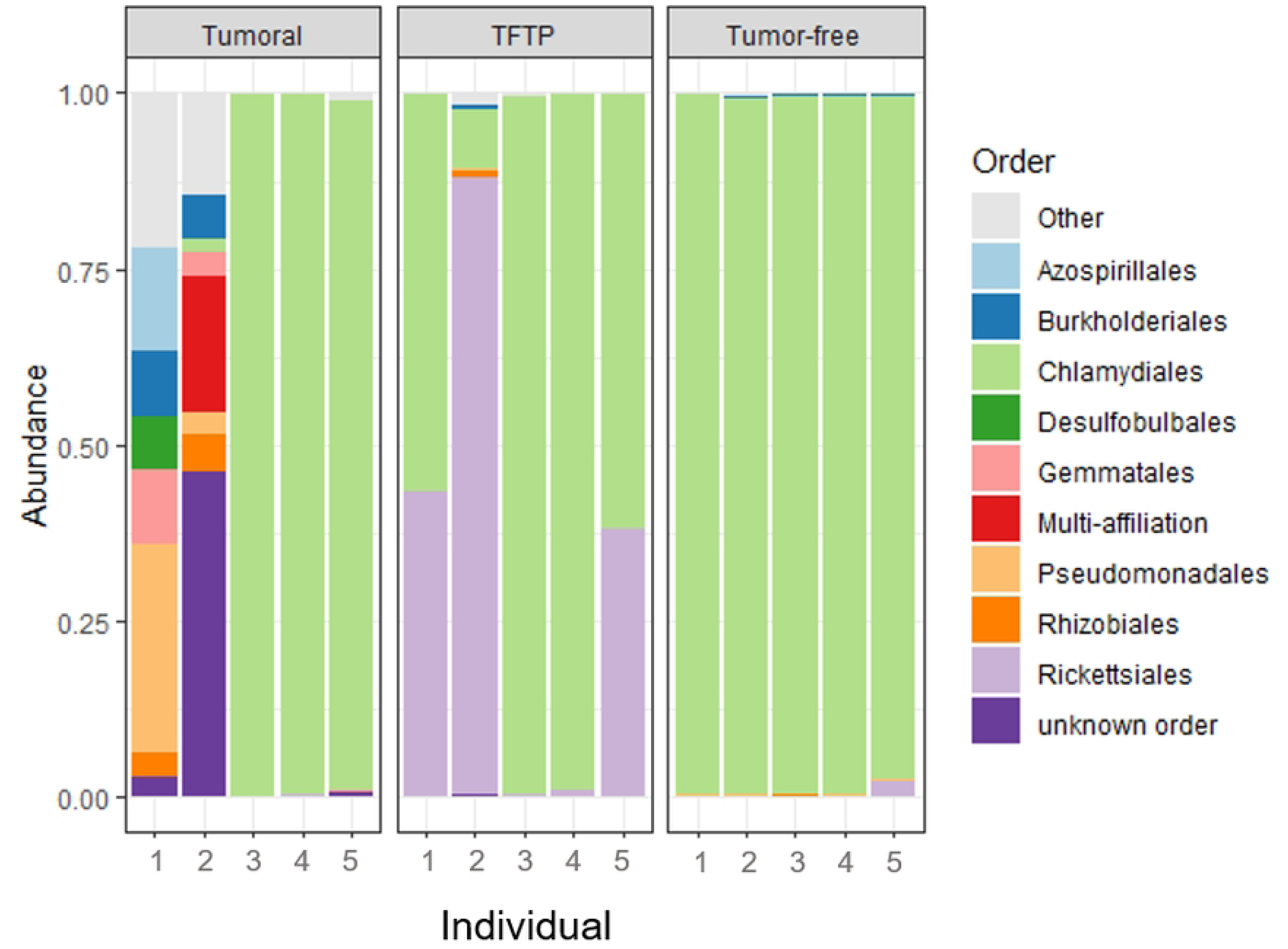
Abundance of the ten most represented bacterial orders in the microbiome of each individual analyzed in each status.

## Discussion

As transmissible cancers are most often detected once they have already spread within host populations, the conditions associated with their emergence remain poorly understood. The cnidarian *H. oligactis* is emerging as a particularly promising model to investigate the evolution of transmissible tumors because it can develop spontaneous tumors relatively easily in the lab [23], and one lab strain harbors a transmissible tumor [24]. In this experimental study, we succeeded for the first time to observe the evolution of a transmissible tumor in the hydra. We have also shown that the rate of transmission of spontaneous tumors can increase over time; namely, 4 times in just 5 generations of selection. Finally, it is remarkable to note that this 5th generation of hydras carrying transmissible tumors is already showing changes in its life history traits compared with its healthy counterparts.

First, this study confirms that *H. oligactis* is a species that easily develops spontaneous tumors in the lab [23]. In line with the tumors described by Boutry et al [23], those from this study (that are coming from hydras collected in the same sampling area) also involve abnormal proliferation of large interstitial stem cells. However, in contrast to the tumors described by Rathje et al [25], these tumors do not appear to be associated with a specific microbiota (since the tumoral and tumor-free individuals from the same strain did not differ from each other in microbiota composition). The reason why *H. oligactis* of the St Petersburg strain, unlike these hydras or those of the *Pelmatohydra robusta* strain (see [25]), diverge in their reliance on a particular microbiome for tumor initiation and maintenance needs to be elucidated by further studies.

Our findings regarding the occurrence of tumors in the descendants of individuals with tumors are *prima facie* consistent with three potential scenarios. One possibility is that these tumors are also spontaneous (i.e., not vertically transmitted from their parents). A second hypothesis suggests parental hydras transmit to their offspring a genetic vulnerability that predisposes them to tumor formation under some conditions. The final possibility is that the tumor itself is transmitted. We favor this last scenario for several reasons. Firstly, the observation that the rate of tumor development in descendants of hydra with tumors is higher than the one of tumor-free hydra reared under the same conditions, implies that these tumors are not spontaneous. In addition, the fact this rate increases over time reinforces this statement. Secondly, the fact that the offspring of TFTP, which share the same genetics as the tumor-bearing hydra but remain tumor-free, do not develop as many tumors as the offspring of tumoral parents, indicates that we cannot attribute the issue to a genetic predisposition for tumor development. Therefore, the most likely explanation for these results is that the tumors observed in the offspring of tumoral parents are transmissible, like those observed in the St Petersburg strain.

Concerning the transmissibility, our findings remarkably indicated that some spontaneous tumors can be immediately transmissible, even if this was not systematic. However, since we lack information on whether the wild individuals used to establish the different strains had different genotypes or were clones when sampled, we cannot draw definitive conclusions regarding the precise proportion of spontaneous tumors that are directly transmissible. To assess the risk of pseudo-replication in this context, further studies would be necessary to estimate the number of genetically distinct hydras in field samples, as well as the probability of two genetically identical hydra both developing spontaneous, transmissible tumors. In addition to be sometimes immediate, tumor transmissibility can be artificially selected across generations. It is important to highlight that if an individual did not develop a tumor within two months, the follow-up observations were terminated. This interruption occurred without knowing whether tumors would later develop or if the individuals had the potential to produce buds that could develop tumors. In both scenarios, this could result in an underestimation of the transmission rate. Despite the fact that tumors have acquired the ability to be transmitted, this ability appears to be fragile, as it is particularly vulnerable to environmental conditions such as the availability of food. For instance, we observed a reduction in the rate of tumor transmission from 90% to 50% in one generation between the end of the experimental selection of tumor transmissibility (5^th^ generation) and the start of the analysis of life history traits. This change coincides with a modification in the feeding protocols between the two experiments, leading to reduced food availability during the measurement of life history traits. This observation is consistent with a previous study [22] that demonstrated a close link between diet and tumoral development in *H. oligactis*. The propensity to develop tumors, as well as the rate at which they develop, is positively correlated with the amount of food. Thus, this study suggests that the ability of spontaneous tumors that acquire transmissibility to maintain it, is also dependent on food intake.

In contrast to the first identified transmissible tumor in hydra (St. Petersburg strain), which exhibits a constant transmission rate (71% at two weeks when asymptomatic and 71% at 5 weeks when symptomatic) [27], our current study on the MT40 strain reveals that the tumor is twice as likely to be transmitted when the host has entered the pathological phase (i.e., developed a tumor) compared to the prepathological phase (25% at two weeks, versus 50% from the fifth week). Several non-exclusive scenarios may account for this disparity. Firstly, during the tumor transmission selection protocol, only buds born during the pathological phase of their parents were surveyed, potentially biasing the selection towards transmission during the pathological phase and resulting in a lower transmission rate in the prepathological phase. A second explanation is that the tumors from the St. Petersburg strain had more time (i.e. 15 years in mass culture) to optimize their transmission, thereby improving it even during the early stages of the tumorigenesis, i.e. during the pre-pathologic stage. This hypothesis would further support the conclusion that transmissibility is a selectable trait. Another explanation could be attributed to the distinct etiology of the tumor. In the St. Petersburg strain, tumor development is induced and persists only in the presence of a specific microbiome [25], absent in the MT40 strain. Our results could be explained if we assume that microbiome transmission remains constant regardless of the age of the hydra, leading to similar tumor development thereafter.

However, when tumor development depends on the initial tumor cells’ number transmitted rather than the microbiome (i.e. MT40), it is expected that symptomatic individuals could be more likely to infect their offspring than those in the pre-pathological stage.

Regarding life-history traits, the tumoral hydra of the MT40 strain exhibits, compared to a tumor-free population, (i) a similar first bud date, (ii) an initial increase in asexual reproductive effort only preceding tumor development, followed by a substantial decrease in budding rate thereafter, (iii) higher mortality in buds produced after the appearance of tumors, and (iv) a comparable survival time. The tumor appears to exert a detrimental influence on hydra budding immediately after the development of external tumoral manifestations, without diminishing overall survival. This could suggest a castration-like phenomenon akin to certain host-parasite systems [39], ultimately leading to a decline in fitness. In conjunction with the lower transmissibility of the tumor in the prepathological phase (i.e., before tumor appearance), these modifications suggest an adjustment of life-history traits of the host to offset the tumor’s costs by producing more buds when they are more likely to survive and remain tumor-free. Interestingly, Boutry et al. [27] also found that tumoral hydras from the St. Petersburg strain adjust their life history traits. However, what is remarkable here is that such adjustments can appear from the 5th generation, i.e. in only a few months. Further research is needed to specify when hydras bearing transmissible tumors start to adjust their life-history traits.

This study has enabled the evaluation, for the first time, of transgenerational effects of tumors on their host by investigating life-history traits in TFTP. These polyps have similar characteristics to those of tumoral polyps during the prepathological phase, but their asexual reproduction rate does not diminish after the fifth week as in tumoral polyps. Consequently, it seems that hydra that do not inherit tumors from their tumoral parents still inherit some tumoral factors that induce an early increase in budding, and this increase is remarkably not subsequently hindered by the presence of tumors. This result, for the initial nine weeks, in a superior asexual fitness compared to both tumoral and tumor-free polyps. Although we have not been able to measure trade-offs in TFPT, we cannot rule out that more in-depth studies would detect them. This could manifest as a reduction in budding rates occurring later (beyond nine weeks) or modifications in sensitivity to biotic and abiotic stress, for instance. Therefore, further research is necessary to comprehensively evaluate the transgenerational effects of tumors on overall fitness.

In the Perfect Storm theory initially proposed by Ujvari et al. [15], and further developed by Tissot et al. [16], two critical barriers have been proposed to explain the evolution of transmissible cancers. The first barrier pertains to the acquisition of transmissibility by tumor cells, while the second relates to the capacity to spread in host populations. The results of the present study suggest, at least in a hydra model, that the first barrier does not appear to be a major obstacle since a significant proportion of spontaneous tumors are immediately transmissible. Indirectly, these results support the idea that the scarcity of transmissible cancers in ecosystems is likely more attributed to the lack of suitable ecological conditions for their spread within populations, i.e. the second barrier (e.g., the presence of predators that would rapidly eliminate diseased individuals, or the sensitivity of tumors to environmental stress – such as food availability). If this conclusion is confirmed in the future, it is crucial to consider these aspects in the study of ecosystems disturbed by human activities, as they could potentially modify the conditions that favor the spread of transmissible cancers (see [40]).

## Material and methods

### Sampling of wild hydra and tumor induction

The hydra used in this study were collected from the Montaud lake in France (43°44’52”N; 3°59’23”E) on May 2^nd^, 2022. Fifty hydras were individually maintained at 18°C in cell culture plates (12-well plates, 1.5 mL/well, Thermo Scientific) filled with Volvic© water under a photoperiod of 12h. To ensure a high rate of budding [26], tumor development [22], and thus the chances of tumor transmission, some of the polyps were fed *ad libitum* five times a week with nauplii of *Artemia salina,* obtained as described in [27]. The wells were cleaned eight hours after feeding by removing the leftover artemia and refilled with Volvic© water. The development of tumor in this initial generation of hydras and in their descendants was characterized with the help of the scale used by Tissot et al. [24]. Hydras were considered as tumoral when they showed at least one medium deformation of their body (Figure 5). An additional 100 polyps, placed in mass culture, were fed *ad libitum* only three times a week to restrict the occurrence of spontaneous tumors, consequently serving as the initial (F0) control population of tumor-free hydras (CTRL).

**Figure 5.**
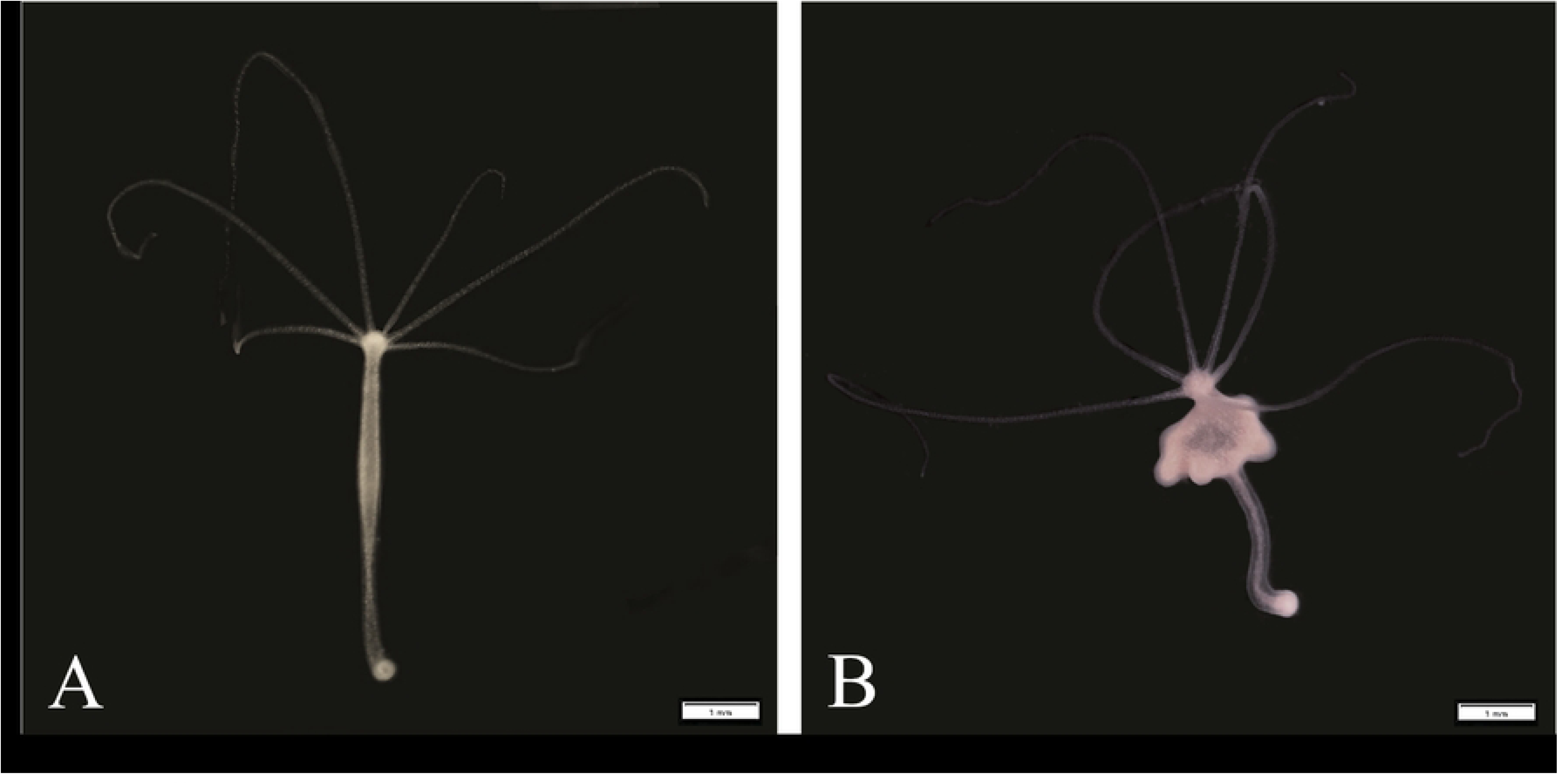
Phenotype of tumor-free and tumoral hydras from the laboratory population established with individuals sampled from Montaud lake. (A) Tumor-free hydra from the control population: the body is long and thin. (B) Hydra from the tumoral strain presenting numerous masses thickening the body. A trinocular magnifier was used to take the pictures, scale bar: 1 mm.

### Selection for tumor transmissibility

To estimate initial tumor transmission rate, we isolated 19 individuals (F0) from the hydras fed five times a week as soon as they have developed spontaneous tumors, and collected all the buds that they subsequently produced until their death. These buds (F1) were placed individually under the same conditions as their parents, and their health status (i.e. development of tumor or not) over time was monitored. When a tumor develops in the F1 cohort, it may be due to transmission from the F0 parents, or to the spontaneous development of a tumor in an F1 individual. In an attempt to control for the rate of spontaneous tumor emergence, we simultaneously isolated buds (F1 CTRL) from the initial population of tumor-free hydras (F0 CTRL) and placed them individually under the same rearing conditions for two months, with five feedings per week. At the end of this period, their health status (i.e. tumor-free, tumoral, or dead) was recorded. In buds from F0 tumoral parents, if tumors appeared within two months, the date of tumor appearance was recorded. Then, all buds produced after that date until the individual’s death were isolated and surveyed in the same manner as their parents. If no tumors appeared within two months, the survey was stopped, and the health status was recorded as “tumor-free”. If an individual died during the experimentation, the associated date was also recorded, and the status was marked as “dead”. This process was repeated during 8 months in order to obtain a total of 4 generations and was then stopped for logistic reasons. Figure 6 summarizes the protocol used to select for tumor transmissibility from hydras with spontaneous tumors.

**Figure 6.**
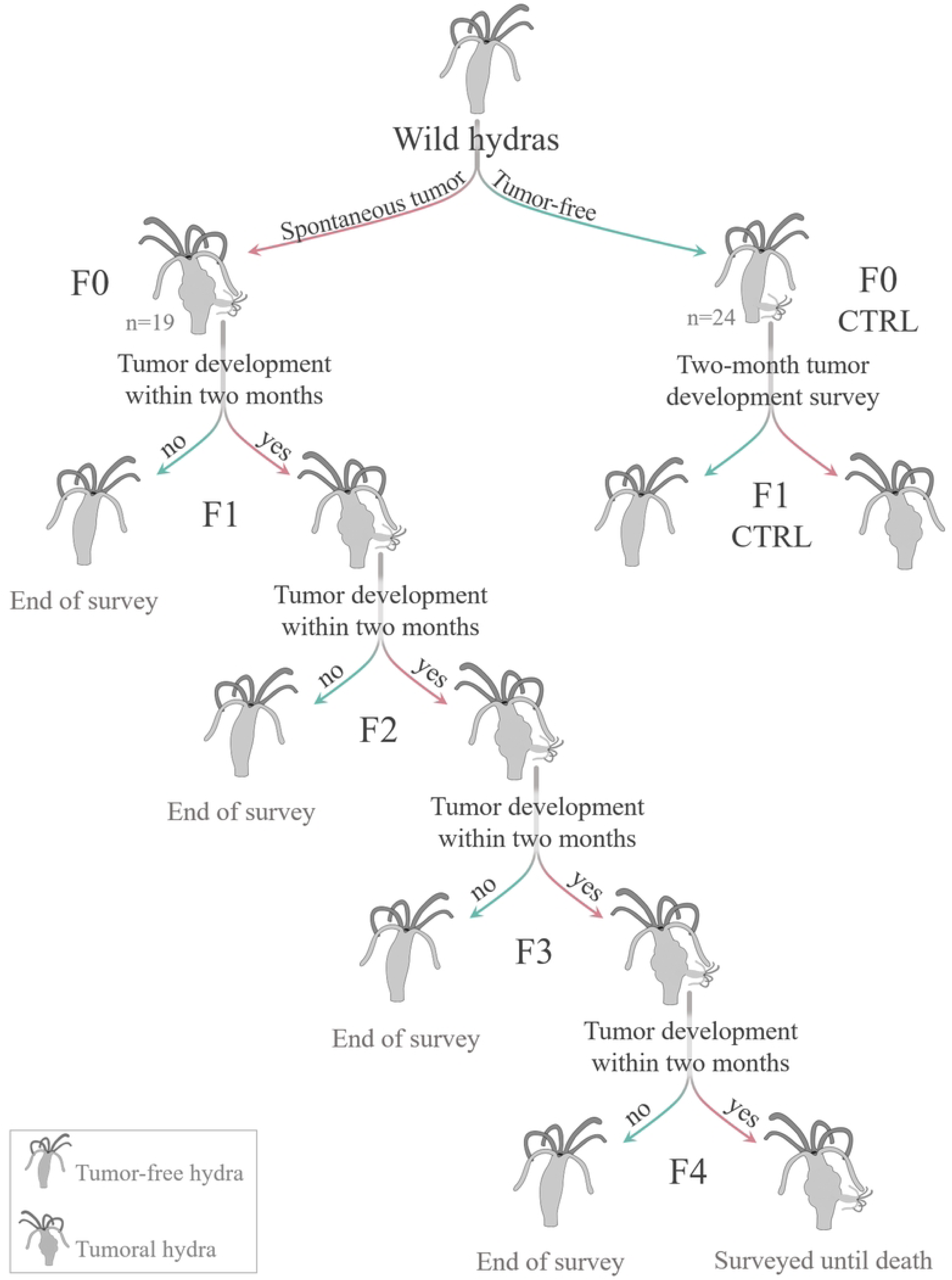
Illustration of the protocol for selection on tumor transmissibility.

### Characterization of a selected hydra strain with transmissible tumors

#### Life-history traits

The fifth generation of a tumoral hydra strain, namely MT40, obtained from the selection experiment described above, was used to quantify its life history traits. We used this strain because it had the highest budding rate, allowing us to obtain enough individuals to carry out this type of analysis. The tumor-free polyps from the same initial population were used as controls. To estimate the asexual reproductive effort, we monitored 46 buds from tumor-free hydras and 63 from the MT40 tumoral strain over six months. Of the 63 hydras derived from the tumoral strain, 26 have developed tumors and were identified as tumoral, while the 37 that have not developed tumors will henceforth be referred to as TFTP (Tumor-Free hydra from Tumoral Parent). Figure 7 summarizes the protocol used to measure life history traits on the strain of interest and its control. This analysis involved tracking the age at first budding, the weekly bud count for 9 weeks, bud survival at two months, tumor transmission rate (i.e., the number of infected buds out of the total bud count), and the survival time (in days). To measure bud survival and tumor transmission rate, firstly during weeks 0 to 4, we isolated 84 buds from tumoral hydras during their prepathological phase (i.e. before tumor appearance, see introduction), 42 buds from TFTP, and 29 buds from tumor-free hydras as control. Then, from the same polyps, during weeks 5 to 9 corresponding to the pathological phase for tumoral hydras (i.e. after tumor appearance), we isolated 40 buds from tumoral polyps, 68 from TFTP, and 35 from tumor-free hydras as control. All the buds isolated in the prepathological and in the pathological phase were surveyed for two months, at the end of which their status was evaluated (i.e. tumoral, tumor-free or dead).

**Figure 7.**
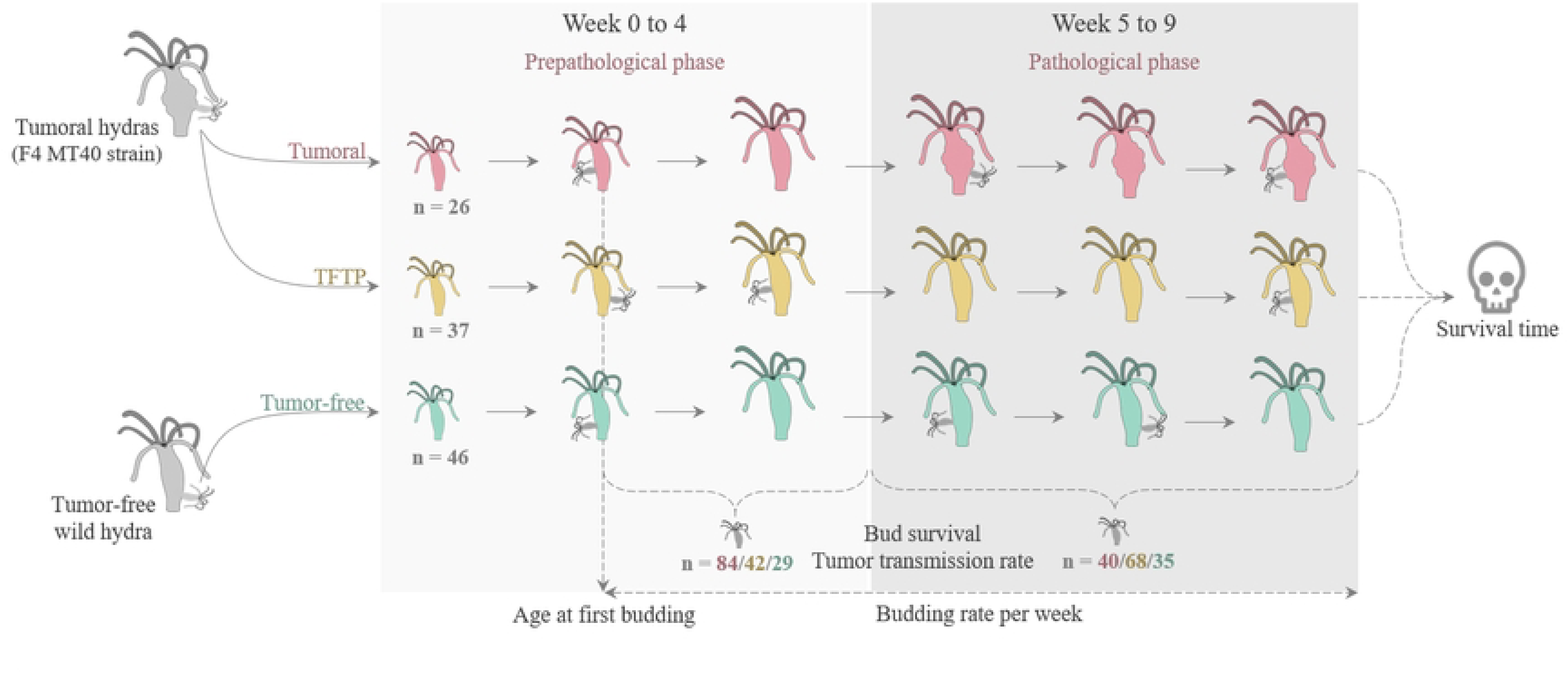
Graphical summary of the protocol for life-history trait measurements.

#### Microbiome and cell type

To characterize the etiology of the tumors, we conducted a microbiome and cell type analysis. This aimed to determine (i) whether they are associated with a specific microbiome, as observed in the transmissible hydra tumors described by Domazet-Lošo et al [24], and (ii) whether the cell type forming the tumor is similar to that reported by Boutry et al [23] in the same hydra population.

For the microbiome analysis, we utilized 5 tumor-free hydras from the control population, 5 tumoral hydras, and 5 TFTP. They were first washed in three baths of sterile water before being placed at –20°C for storage. DNA was extracted with the Qiagen Blood & Tissue Kit according to the manufacturer’s instructions, and DNA concentration was measured using a Qubit 2.0 fluorometer. The primers 5’GTGCCAGCMGCCGCGGTAA and 3’GGATTAGAWACCCBDGTAGTCC were used to target the V4 region of the 16S gene during the PCR. The products were then sent to the genomic platform (Genseq, Montpellier University) for Illumina sequencing.

For the cell type analysis, two individuals from each status, namely tumor-free (as control), tumoral, and tumor-free from tumoral parent were macerated together and 100µL of the solution were spread on gelatin-coated microscope slides according to the procedure detailed in [28]. Once dry, the slides were observed by phase contrast under a microscope with a 40x objective, and the number of epithelial, small interstitial stem cells, and large interstitial stem cells were counted. Two individuals were macerated for each status.

### Data analysis

The analyses presented here were performed with the R software (version 4.2.2) [29] and the graphical representations were realized with the “GGplot2” package [30]. The sequencing data for the microbiome were checked for their quality and processed *via* the FROGS analysis pipeline developed by the GenoToul genomic platform in the Galaxy interface [31].

#### Selection for tumor transmissibility

To analyze (i) the proportion of tumoral hydras in the first generation (F1) according to the parental status (i.e. tumoral or tumor-free) and (ii) the tumor transmission rate according to generation and bud order (i.e. the ratio of bud rank on total bud number per parent), we used generalized linear mixed-effects models (GLMMs) from the package “glmmTMB” [32], as the response variables here were non-normally distributed (i.e. binary or count data). For the proportion of hydras developing tumors in F1, a random intercept effect of the birth date was added to consider the possible temporal variability, and a binomial distribution was used. Concerning tumor transmission rate, a random intercept effect of the founder individual (i.e. F0 strain) was added to consider possible variability between strains, and a binomial distribution was applied. In this analysis, buds without siblings were excluded (constituting 3 cases out of 280) to specifically investigate the impact of bud order on tumoral transmission. According to Zuur et al. [33], model selection was made first on the random effect and then on the fixed effect, based on the weight of the corrected Akaike information criterion (AICc) obtained with the package “MuMIn” [34]. Then, the fitting of the model obtained was checked with “DHARMa” package [35] which performs a Kolmogorov-Smirnov, outlier, and overdispersion test on model residuals. The detail of the variable types (i.e. in this case, continuous, date or factor variables) as well as all the models constructed and the AICc weight associated is presented in the S1 Table in the supporting information.

#### Life-history traits

To analyze (i) first bud date according to status, (ii) budding rate according to status and phase (i.e. from the first bud date to the week 4 and weeks 5 to 9, corresponding to the pre-pathological and pathological phases of tumoral hydras, respectively), (iii) bud survival according to the status and the phase, and (iv) tumor transmission rate according to status and phase, we used used GLMMs. Concerning the age at first budding and the budding rate analysis, a random intercept effect of birthday was added to control variability linked respectively to different batches, and only for budding rate the random effect of the individual was also added to consider the repeated measurements on the same individuals. A random effect of the parent’s identity was added to analyze bud survival and tumor transmission rate, as buds collected in both phases come from the same parents. For the analysis of the first two traits, a negative binomial distribution was used as the age at first budding counts the number of failures (i.e. days) before the success (i.e. the first bud production) and the budding rate is too low to use a Poisson distribution (i.e. the variable is underdispersed). For the last two variables, a binomial distribution was used as they are both binary. The model selection and validation are identical to the previous section. With regard to the age at first budding, an outlier was removed from the dataset (58 days vs. an average of 17) after being detected by the outlier test during verification of the established model, which prevented a good fit. Concerning the analysis of bud survival and tumor transmission rate, as only one control individual developed a spontaneous tumor, it was removed from the data set in order to test the presence of an interaction between status and phase. The analysis of tumor transmission rate was performed only on buds from tumoral hydras and TFTP, as no tumor appearance was observed on buds from control hydras.

For the analysis of the survival time according to the status, survival regressions were used with the exponential distribution, as the instantaneous risk was constant (verification by plotting the survival curve of individuals). The selection model was also based on the AICc weight. The details of variable forms, as well as all the models tested and their AICc weights, are presented in the S2 Table in the supporting information.

#### Cell type and Microbiome

Only the ratio between interstitial stem cells and epithelial cells has been calculated on the data collected after maceration (see [23]).

The bacterial feature table obtained from the FROGS pipeline was used to compare treatment groups in terms of alpha and beta diversity. For alpha diversity, we calculated the chao1 index as a proxy for bacterial species richness and used Kruskal-Wallis test to compare treatment groups. For beta diversity, we calculated distance between samples using the Bray-Curtis index (which takes bacterial abundance into account) and the Jaccard index (which takes only presence-absence data into account). These distance matrices were subsequently compared using Adonis with N=999 permutations and Pairwise Adonis to test the statistical null hypothesis that the microbiome composition of treatment groups is not differentiated. Microbiome analyzes were performed with the *microbiome* [36], *vegan* [37] and *pairwiseAdonis* [38] R packages.

## Author contributions

The original idea for the article and the design of the study came from S.T with the support of F.T. S.T and J.M. collected the data on hydra. L.B and J.T performed microbiome analyses and S.T performed the cell type analysis and statistical analysis supported by A.M.D. S.T wrote the manuscript with the support of F.T, B.U, A.M.D, A.M.N, J.B, J.T, J.M, and B.R.

## Data availability statement

Scripts and data associated are provided in supplementary information. The 16S rRNA raw sequence files for this study have been deposited in FASTQ format and can be found in the Sequence Read Archive from NCBI (BioProject: PRJNA1072327).

## Acknowledgements

This work was carried out within the framework of the Camargue Health-Environment “Zone Atelier” (ZACAM) of the “Long-Term Socio-Ecological Research network” (LTSER-RZA), funded by the Ecology & Environment of the French National Center for Scientific Research (EE-CNRS).

## Supporting information

**S1 File. Data and scripts used in this study.** This file contains 7 data sets used in 4 scripts: “microbiome” for analysis of microbiome information, “Script transmission” for analysis of transmission selection, and “THV-script” for analysis of life history characteristics, with the exception of bud survival and tumor transmission rate, which are present in “THV-bud_becoming”.

**S1 Table. The formulas of the different models constructed and the associated value of AICc’s weight.** A double line separates the selection of random effects from fixed effects, and shaded boxes indicate that the model has not converged. “Status of the parent”, “Status of hydra in the first generation”, “Status of hydra”, and “ID founder individual” are factor variables. “Birthday” is a date variable. “Bud order”, and “Generation” are continuous variables.

**S2 Table. The formulas of the different models constructed and the associated value of AICc (for model with random effect) or AIC’s weight (for model without random effect).** A double line separates the selection of random effects from fixed effects, and shaded boxes indicate that the model has not converged. “Status of the parent”, “Status”, “Phase”, “Tumor transmission rate” and “ID” are factor variables. “Birthday” is a date variable. “Age at first budding”, “Budding rate”, “Bud survival”, and “Survival time” are continuous variables.

**S1 Figure. Graphical representations of alpha diversity (A), principal coordinate analysis (PCoA) of the Bray-Curtis index (B), and principal coordinate analysis of the Jaccard index (C), all according to the status.**

## References

[1] C. A. Aktipis et R. M. Nesse, «Evolutionary foundations for cancer biology», Evol. Appl., vol. 6, n° 1, p. 144–159, 2013, doi: 10.1111/eva.12034.

[2] B. Ujvari, B. Roche, et F. Thomas, Ecology and Evolution of Cancer. Academic Press, 2017.

[3] A. M. Dujon, G. Bramwell, B. Roche, F. Thomas, et B. Ujvari, «Transmissible cancers in mammals and bivalves: How many examples are there?», BioEssays, vol. 43, n° 3, p. 2000222, 2021, doi: 10.1002/bies.202000222.

[4] C. A. Aktipis et al., «Cancer across the tree of life: cooperation and cheating in multicellularity», Philos. Trans. R. Soc. B Biol. Sci., vol. 370, n° 1673, p. 20140219, juill. 2015, doi: 10.1098/rstb.2014.0219.

[5] R. Pye et al., «Demonstration of immune responses against devil facial tumour disease in wild Tasmanian devils», Biol. Lett., vol. 12, n° 10, p. 20160553, oct. 2016, doi: 10.1098/rsbl.2016.0553.

[6] B. Ganguly, U. Das, et A. K. Das, «Canine transmissible venereal tumour: a review», Vet. Comp. Oncol., vol. 14, n° 1, p. 1–12, mars 2016, doi: 10.1111/vco.12060.

[7] D. Garcia-Souto et al., «Mitochondrial genome sequencing of marine leukaemias reveals cancer contagion between clam species in the Seas of Southern Europe», eLife, vol. 11, p. e66946, janv. 2022, doi: 10.7554/eLife.66946.

[8] M. J. Metzger, C. Reinisch, J. Sherry, et S. P. Goff, «Horizontal transmission of clonal cancer cells causes leukemia in soft-shell clams», Cell, vol. 161, n° 2, p. 255–263, avr. 2015, doi: 10.1016/j.cell.2015.02.042.

[9] M. J. Metzger et al., «Widespread transmission of independent cancer lineages within multiple bivalve species», Nature, vol. 534, n°7609, p. 705–709, juin 2016, doi: 10.1038/nature18599.

[10] M. A. Yonemitsu et al., «A single clonal lineage of transmissible cancer identified in two marine mussel species in South America and Europe», eLife, vol. 8, p. e47788, nov. 2019, doi: 10.7554/eLife.47788.

[11] M. Hammel et al., «Marine transmissible cancer navigates urbanized waters, threatening spillover», Proc. R. Soc. B Biol. Sci., vol. 291, n° 2017, p. 20232541, févr. 2024, doi: 10.1098/rspb.2023.2541.

[12] M. R. Stammnitz et al., «The evolution of two transmissible cancers in Tasmanian devils», Science, vol. 380, n°6642, p. 283–293, avr. 2023, doi: 10.1126/science.abq6453.

[13] R. Hamede et al., «The tumour is in the detail: Local phylogenetic, population and epidemiological dynamics of a transmissible cancer in Tasmanian devils», Evol. Appl., vol. 16, n° 7, p. 1316–1327, 2023, doi: 10.1111/eva.13569.

[14] D. Hanahan et R. A. Weinberg, «Hallmarks of Cancer: The Next Generation», Cell, vol. 144, n°5, p. 646–674, mars 2011, doi: 10.1016/j.cell.2011.02.013.

[15] B. Ujvari, R. A. Gatenby, et F. Thomas, «Transmissible cancers, are they more common than thought?», Evol. Appl., vol. 9, n° 5, p. 633–634, 2016, doi: 10.1111/eva.12372.

[16] S. Tissot et al., «Transmissible Cancer Evolution: The Under-Estimated Role of Environmental Factors in the “Perfect Storm” Theory», Pathogens, vol. 11, n°2, Art. n° 2, févr. 2022, doi: 10.3390/pathogens11020241.

[17] E. a. V. Burioli et al., «Traits of a mussel transmissible cancer are reminiscent of a parasitic life style», Sci. Rep., vol. 11, n° 1, Art. n° 1, déc. 2021, doi: 10.1038/s41598-021-03598-w.

[18] C. E. Hawkins et al., «Emerging disease and population decline of an island endemic, the Tasmanian devil Sarcophilus harrisii», Biol. Conserv., vol. 131, n° 2, p. 307–324, août 2006, doi: 10.1016/j.biocon.2006.04.010.

[19] C. Hogg, S. Fox, D. Pemberton, et K. Belov, Saving the Tasmanian Devil: Recovery through Science-based Management. Csiro Publishing, 2019.

[20] T. Hollings, M. Jones, N. Mooney, et H. Mccallum, «Trophic Cascades Following the Disease-Induced Decline of an Apex Predator, the Tasmanian Devil», Conserv. Biol., vol. 28, n° 1, p. 63–75, 2014, doi: 10.1111/cobi.12152.

[21] M. Vittecoq et al., «Cancer: a missing link in ecosystem functioning?», Trends Ecol. Evol., vol. 28, n° 11, p. 628–635, nov. 2013, doi: 10.1016/j.tree.2013.07.005.

[22] S. Tissot et al., «The impact of food availability on tumorigenesis is evolutionarily conserved», Sci. Rep., vol. 13, n° 1, Art. n° 1, nov. 2023, doi: 10.1038/s41598-023-46896-1.

[23] J. Boutry et al., «Spontaneously occurring tumors in different wild-derived strains of hydra», Sci. Rep., vol. 13, n° 1, Art. n° 1, mai 2023, doi: 10.1038/s41598-023-34656-0.

[24] T. Domazet-Lošo et al., «Naturally occurring tumours in the basal metazoan Hydra», Nat. Commun., vol. 5, n° 1, Art. n° 1, juin 2014, doi: 10.1038/ncomms5222.

[25] K. Rathje, B. Mortzfeld, M. P. Hoeppner, J. Taubenheim, T. C. G. Bosch, et A. Klimovich, «Dynamic interactions within the host-associated microbiota cause tumor formation in the basal metazoan Hydra», PLOS Pathog., vol. 16, n° 3, p. e1008375, mars 2020, doi: 10.1371/journal.ppat.1008375.

[26] S. Gurkewitz, M. Chow, et R. D. Campbell, «Hydra size and budding rate: influence of feeding», Int. J. Invertebr. Reprod., vol. 2, n° 3, p. 199–201, janv. 1980, doi: 10.1080/01651269.1980.10553355.

[27] J. Boutry et al., «Tumors alter life history traits in the freshwater cnidarian, Hydra oligactis», iScience, vol. 25, n° 10, Art. n° 10, oct. 2022, doi: 10.1016/j.isci.2022.105034.

[28] C. N. David, «Dissociating Hydra Tissue into Single Cells by the Maceration Technique », in *Hydra: Research Methods*, H. M. Lenhoff, Éd., Boston, MA: Springer US, 1983, p. 153–156. doi: 10.1007/978-1-4757-0596-6_21.

[29] R Development Core Team, « R: a language and environment for statistical computing ». R Foundation for Statistical Computing, Vienna, Austria, 2020.

[30] H. Wickham et al., « ggplot2: Create Elegant Data Visualisations Using the Grammar of Graphics ». 10 février 2023. Consulté le: 13 février 2023. [En ligne]. Disponible sur: https://CRAN.R-project.org/package=ggplot2

[31] M. Ranchou-Peyruse et al., «A deep continental aquifer downhole sampler for microbiological studies», Front. Microbiol., vol. 13, 2023, Consulté le: 15 janvier 2024. [En ligne]. Disponible sur: https://www.frontiersin.org/articles/10.3389/fmicb.2022.1012400

[32] M. Brooks et al., « glmmTMB: Generalized Linear Mixed Models using Template Model Builder ». 12 juillet 2022. Consulté le: 29 octobre 2022. [En ligne]. Disponible sur: https://CRAN.R-project.org/package=glmmTMB

[33] A. F. Zuur, E. N. Ieno, N. Walker, A. A. Saveliev, et G. M. Smith, Mixed effects models and extensions in ecology with R. in Statistics for Biology and Health. New York, NY: Springer, 2009. doi: 10.1007/978-0-387-87458-6.

[34] K. Bartoń, « MuMIn: Multi-Model Inference ». 1 septembre 2022. Consulté le: 29 octobre 2022. [En ligne]. Disponible sur: https://CRAN.R-project.org/package=MuMIn

[35] F. Hartig et L. Lohse, « DHARMa: Residual Diagnostics for Hierarchical (Multi-Level / Mixed) Regression Models ». 8 septembre 2022. Consulté le: 29 octobre 2022. [En ligne]. Disponible sur: https://CRAN.R-project.org/package=DHARMa

[36] L. Lahti, S. Shetty, et et al, « Tools for microbiome analysis in R ». [En ligne]. Disponible sur: http://microbiome.github.com/microbiome.

[37] J. Oksanen et al., « vegan: Community Ecology Package ». 11 octobre 2022. Consulté le: 12 février 2024. [En ligne]. Disponible sur: https://cran.r-project.org/web/packages/vegan/index.html

[38] P. M. Arbizu, « pmartinezarbizu/pairwiseAdonis ». 2017. Consulté le: 12 février 2024. [En ligne]. Disponible sur: https://github.com/pmartinezarbizu/pairwiseAdonis

[39] K. D. Lafferty et A. M. Kuris, «Parasitic castration: the evolution and ecology of body snatchers», Trends Parasitol., vol. 25, n° 12, p. 564–572, déc. 2009, doi: 10.1016/j.pt.2009.09.003.

[40] A. M. Dujon et al., «On the need for integrating cancer into the One Health perspective», Evol. Appl., vol. 14, n° 11, p. 2571–2575, 2021, doi: 10.1111/eva.13303.

